# Cross-source proteogenomic profiling of human microglia reveals a conserved core membrane protein and HERV signature

**DOI:** 10.64898/2026.09.25.754430

**Authors:** Hoi Tong Wong, Charles Gleason, Andrew Boylan, Michelle Adler, Roman Kosoy, John F. Fullard, Elizabeth Levendosky, Gabriel E. Hoffman, Henrik Molina, Panos Roussos, Dusan Bogunovic, Viviana Simon, Lubbertus C.F. Mulder

**Affiliations:** Department of Microbiology, Icahn School of Medicine at Mount Sinai, New York, NY, USA; Center for Vaccine Research and Pandemic Preparedness, Icahn School of Medicine at Mount Sinai, New York, NY, USA; Department of Pediatrics, Columbia University, New York, NY, USA; Marc and Jennifer Lipschultz Precision Immunology Institute, Icahn School of Medicine at Mount Sinai, New York, NY 10029, USA; Department of Psychiatry, Icahn School of Medicine at Mount Sinai, New York, NY, USA; Center for Disease Neurogenomics, Icahn School of Medicine at Mount Sinai, New York, NY, USA; Department of Genetics and Genomic Sciences, Icahn School of Medicine at Mount Sinai, New York, NY, USA; Friedman Brain Institute, Icahn School of Medicine at Mount Sinai, New York, NY, USA; Proteomics Resource Center, Rockefeller University, New York, NY, USA; Mental Illness Research, Education and Clinical Center VISN2, James J. Peters VA Medical Center, Bronx, NY, USA; Center for Precision Medicine and Translational Therapeutics, James J. Peters VA Medical Center, Bronx, NY, USA; Center for Genetic Errors of Immunity, Columbia University, New York, NY, USA; The Global Health and Emerging Pathogen Institute, Icahn School of Medicine at Mount Sinai, New York, NY, USA; Division of Infectious Diseases, Department of Medicine, Icahn School of Medicine at Mount Sinai, New York, NY, USA; Department of Pathology, Molecular and Cell based Medicine, Icahn School of Medicine at Mount Sinai, New York, NY, USA

**Author notes:** Correspondence to Lubbertus C.F. Mulder.

**Keywords:** HERV, human endogenous retroviruses, monocyte-derived microglia, brain-derived microglia, iPSC-derived microglia, RNA-seq, mass spectrometry, neurodegenerative disease

## Abstract

Microglia are essential immune cells of the brain parenchyma which are responsible for maintaining homeostasis and responding to brain injury. When activated, they induce immune signaling and become phagocytic. In the context of neurodegenerative diseases, they exhibit both pro- and anti-inflammatory properties. Human endogenous retroviruses (HERVs) are the remnants of ancient retroviral germline infections that integrated into the human genome, and their expression has been directly linked to microglial-mediated neurodegenerative diseases. Despite this, HERV expression is not consistently included in analyses characterizing microglia. Investigating the function of microglia across different diseases is critical for elucidating mechanisms of disease progression and for the development of effective therapeutic interventions. Approaches to studying microglia include isolation from post-mortem brain tissues, differentiation from peripheral blood monocytes or differentiation of induced pluripotent stem cells (iPSC). *Ex vivo* brain derived microglia (BdMG) are considered most physiologically relevant, but constraints including the unpredictable timing of availability, labor-intensive protocols, and low yields reduce their practicality. Existing alternatives are iPSC-derived microglia (iMG) and monocyte-derived microglia (MDMi) models that circumvent sample accessibility and scalability issues. Here, we explored the transcriptome and proteome of these three microglia sources and identified a common microglia surface marker expression profile, establishing that MDMi and iMG exhibit similar phenotypes relative to BdMG. We also identified a common HERV expression profile across BdMG, iMG and MDMi, indicating deeper similarities across different sources. Altogether, our findings indicate that given the phenotypic similarities of the 2 models, MDMi represent a practical and reliable model for investigating microglial function with iMG models serving for more molecular based approaches, such as CRISPRa/i.

## Introduction

Microglia are the principal resident myeloid cells of the central nervous system (CNS), accounting for 10–15% of total CNS cells (Lawson et al., 1992). They originate from erythromyeloid precursors in the yolk sac, colonize the developing brain before birth (Ginhoux et al., 2010, Kierdorf et al., 2013), and although they persist into adulthood as a long-lived, self-renewing population (Réu et al., 2017), during life, increasing portions are replaced with microglia originating from myeloid precursors from the bone marrow (Belk et al., 2026). Under homeostatic conditions, microglia carry out continuous immunosurveillance of the brain parenchyma by monitoring neuronal health, tissue integrity, and blood flow (McKinney et al., 2026). During development and throughout adult life they also prune supernumerary synapses (Tay et al., 2017), dispose of apoptotic neurons, cellular debris, and misfolded proteins (McKinney et al., 2026).

Microglia role of in neurodegeneration is context dependent and can be either protective or detrimental. In Alzheimer’s disease (AD), microglia phagocytose and degrade amyloid plaques yet simultaneously promote tau propagation and chronic neuroinflammation (Gao et al., 2023). In multiple sclerosis (MS), they induce demyelination through pro-inflammatory cytokine secretion but also promote remyelination by recruiting oligodendrocyte progenitor cells (Baaklini et al., 2023, van Wageningen et al., 2019). In amyotrophic lateral sclerosis (ALS), microglia clear TDP-43 aggregates during early disease stages, but, when chronically activated, secrete cytotoxic levels of TNF-α, IL-1β, and IL-6, accelerating motor neuron loss (Noh et al., 2025, Bond et al., 2025).

Since microglia occupy a central role across multiple pathologies, biologically relevant human microglia models are essential. Studies that directly compare different sources, including post-mortem brain-derived microglia (BdMG), monocyte-derived microglia (MDMi), iPSC-derived microglia (iMG), and immortalized cell lines such as HMC3, are needed to establish which model best fit a given experimental question (Gumbs et al., 2022, Woolf et al., 2025, Ryan et al., 2017). HMC3 cells are the most accessible of these models, as they are SV40-immortalized and retain some microglial markers, although recent work has demonstrated that they don’t express canonical myeloid markers such as Iba1 (AIF1), CD45, and PU.1, raising questions about their fidelity as a microglia model (Woolf et al., 2025).

iMG models attain transcriptional and functional features of brain microglia with greater fidelity; however, differentiation protocols vary considerably in cytokine cocktails, media composition, and duration, typically requiring multiple weeks of treatment. This hinders yield, scalability, and banking (Teo et al., 2024, Haskell et al., 2026), which is especially constraining for downstream applications devoid of amplification steps, such as mass spectrometry-based proteomics.

MDMi offer a more practical alternative; they are derived from peripheral blood monocytes using cytokine cocktails (often including GM-CSF and IL-34) in 10–14 days, compared to the 40-day multi-step protocols required for iMG. MDMi express microglia markers including P2RY12, TREM2, and APOE, and recapitulate disease-relevant phenotypes (Ryan et al., 2017, Quek et al., 2022). However, like iMGs, MDMi are not derived from yolk sac precursors, and retain monocyte-lineage markers that transcriptionally resemble fetal rather than adult microglia (Bennett et al., 2018). Furthermore, differentiation protocols are non-standardized across laboratories (Sargeant and Fourrier, 2023).

Post-mortem brain tissue remains the gold standard for studying fully differentiated primary microglia. The isolation of microglia from the brain parenchyma is technically demanding and labor-intensive, with conditions that must be meticulously optimized (Mizee et al., 2017, Mizee et al., 2018). The unpredictable timing of availability from brain banks and neurosurgical resections further restricts experimental planning and throughput. Finally, microglia display significant regional heterogeneity, differing in density, morphology, marker expression, and functional capacity across brain regions (Böttcher et al., 2019, Tan et al., 2020).

Human endogenous retroviruses (HERVs) are genomic relics of ancient germ-line retroviral infections that successfully integrated and together encompass 8% of our genome (Lander et al., 2001, Boeke and Stoye, 1997). Expression of HERVs is broadly suppressed in healthy tissue (Turelli et al., 2014, Collins et al., 2015, Cuellar et al., 2017), but limited expression has been shown to be cell type- and tissue-specific at the locus level (Burn et al., 2022, Dopkins and Nixon, 2024) with HERV RNA-transcription widely associated with inflammation (Chuong et al., 2016, Chiappinelli et al., 2015, Wang et al., 2022) .The evolutionary history of HERVs is marked by the accumulation of insertions, deletions, and truncating mutations that progressively abolish their capacity for protein production, with only the most recent proviral elements retaining intact open reading frames suitable for viral polypeptide synthesis (Johnson, 2019). Identification of transcribed HERVs at the locus level remains challenging, and as a result their expression is often not quantified even when RNA-seq data is available. HERVs have been directly linked to microglial-mediated neurodegeneration through the finding that HERV-K (HML-2) envelope RNA is sensed by microglial TLR7/8, triggering a robust pro-inflammatory activation and subsequent neuronal death (Dembny et al., 2020). In MS, the HERV-W envelope protein activates human microglia through TLR4 engagement. The resulting pro-inflammatory phenotype is toxic to oligodendrocytes and inhibits remyelination. This microglial pathway is now being targeted by the investigational therapeutic antibody temelimab (Adler et al., 2024).

GWAS studies have identified neurodegenerative risk variants enriched in microglia-specific gene regulatory elements, including HERV loci which represent shared genomic risk factors in both ALS and MS. Therefore, human microglia represent the cellular interface where endogenous retroviral reactivation converges with genetically determined neuroinflammatory susceptibility (Duarte et al., 2025, Yan et al., 2026).

It has long been recognized that mRNA and protein abundance correlate poorly (Schwanhäusser et al., 2011, Liu et al., 2016, Vogel and Marcotte, 2012, Gygi et al., 1999), and this disconnect is particularly consequential in neurodegeneration. Even in tandem mass spectrometry data derived from postmortem brain tissue, the metabolic and MAPK signaling protein co-expression modules, which were strongly associated with cognitive decline in AD, had no corresponding signal at the transcript level (Johnson et al., 2022). This emphasizes the need of proteomics to capture the functional molecular landscape that transcriptomic analyses cannot resolve. Cell surface protein profiles obtained by mass spectrometry provide a direct, antibody-independent readout of microglial functional identity, enabling rigorous benchmarking across model systems that differ in ontogeny, culture environment, and maturation state (Lloyd et al., 2024).

In this work, we compare the cell-surface transcriptome and proteome of microglia from different sources and extend this comparison to their human endogenous retrovirus (HERV) transcriptome, to provide a resource for selecting the most physiologically relevant model for a given experimental or pathological context.

## Material and methods

### Cell culture

NCCIT germ cell tumor line was purchased from the ATCC (CRL-2073) and cultured in RPMI 20% FBS (GeminiBio, 100-106-500) 100 IU/ml penicillin-streptomycin (Corning, 30-002-CI) and 2 mM L-glutamine (Corning, 25-005-CI). NCCIT cells are an intermediate between seminoma and embryonal carcinoma. HMC3 are immortalized microglia purchased from ATCC (CRL-3304) and cultured in EMEM 10% FBS (GeminiBio, 100-106-500) 100 IU/ml penicillin-streptomycin (Corning, 30-002-CI) and 2 mM L-glutamine (Corning, 25-005-CI). All cells were regularly tested for mycoplasma contamination and kept at 37°C 5% CO2 humidified incubator.

### CD4 T cells and Monocyte isolation and microglia differentiation

Buffy coats were supplied by anonymous healthy donors from by the New York Blood Center (New York Blood Center Component Laboratory) were processed for peripheral blood mononuclear cells (PBMCs) through a density gradient centrifugation using Lymphoprep (STEMCELL technologies, 07861). For CD4 T cell isolation PBMCs underwent magnetic bead negative selection using (CD4+ T cell isolation kit II; Miltenyi Biotec) following manufacturer’s instructions. CD4+ T cells were then cultured in RPMI 1640 supplemented with 10% FCS (Gibco/Thermo Fisher), 100 IU/ml penicillin, 100 μg/ml streptomycin, 0.1 M HEPES, 2 mM l-glutamine, and 20 IU/ml IL-2 for 5 days before downstream analysis. Monocytes were isolated from PBMCs by Percoll gradient (Cytiva, 17089101)(Colotta et al., 1984). Microglia differentiation was obtained by incubating the monocytes in RPMI 1640 (Gibco, 21870-076) with 100 U/ml Penicillin-Streptomycin (Corning, 30-002-CI), 2mM L-glutamine (Corning, 25-005-CI) supplemented with 100ng/ml IL-34 (R&D Systems, 5265-IL), 100ng/ml CCL2 (Biolegend, 571404), 10ng/ml M-CSF (Biolegend, 574804), 10ng/ml GM-CSF (ThermoFisher Scientific, 300-03-20UG) and 10ng/ml β-NGF (R&D systems, 256-GF) cytokines to initiate differentiation into microglia (14). Cells were replenished with cytokines every other day over a period of 9 days, while complete media change was done on the 6^th^ day. All cells were cultured at 37°C in a 5% CO2 humidified incubator until they were ready for extraction.

### Microglia isolation from human brain tissue

For protein analysis, autopsy brain specimens were collected by Mount Sinai Neuropathology Brain Bank & Research CoRE. Microglia isolation was adapted from a previously published protocol (Kosoy et al., 2025). Briefly, fresh sections of occipital lobe and prefrontal cortex were stored in preservation buffer Hibernate A (Transnetyx, HAPR500) or tissue preservation buffer (Milteyni Biotec, 130100008), respectively, at 4°C ≤ 48h before processing. Meninges were manually removed from tissue before processing with the Adult Brain Dissociation Kit (Miltenyi Biotec, 130-107-677) according to manufacturer’s instructions. RNase inhibitors (Takara Bio, 2313B) throughout isolation. Tissues from prefrontal cortex were de-myelinated (Miltenyi Biotec, 130096733) and incubated in the dark with 1:500 antibody cocktail (CD45, clone HI30, BD Pharmingen, 555483; CD11b, clone ICRF44, BD Pharmingen, 560914) for 1h at 4 °C with end-over-end rotation. 4′,6-Diamidino-2-phenylindole (DAPI) (ThermoFisher Scientific, 62248) was added at 1:1,000 dilution; viable (DAPI−) CD45^+^ cells isolated using fluorescence-activated cell sorting and a FACSAria flow cytometer (BD Biosciences). Occipital lobe tissues were selected using CD11b microbeads (Milteyni Biotec, 130-093-634) and magnetic separation according to manufacturer’s instructions. Brain derived microglia RNA-seq data were obtained from previously published dataset (Kosoy et al., 2025). Samples were collected at JJ Peters VA Medical Center NIH Brain and Tissue Repository according to similar parameters of the MSBB–Mount Sinai NIH Neurobiobank cohort (Haroutunian et al., 2009, Wang et al., 2018).

### Stem cell differentiation into microglia

For three of four donors, induced pluripotent stem cells (iPSCs) were generated from PBMCs of healthy male donors (Columbia University IRB AAAV1815) by either New York Stem Cell Foundation (M25HC, M34HC) or Rutgers University Cell and DNA Repository, now known as Infinity BiologiX (DL018) and maintained in mTeSR™ Plus media (StemCell Technologies, 100-0276) on Matrigel-coated plates at 37°C in a 5% CO2 humidified incubator. Cells underwent regular mycoplasma testing and karyotyping. All iPSC differentiation procedures for these three donors were performed by the Stem Cell Core facility at Columbia University Irving Medical Center. iPSCs were differentiated to hematopoietic progenitor cells (HPCs) using the STEMdiff™ Hematopoietic Kit (StemCell Technologies, 05310) with minor modifications to manufacturer’s instructions (using 0.5mM EDTA for dissociation instead of ReLeSR™) and ultimately to microglia (iMG) as described previously (McQuade et al., 2018). Briefly, HPCs were collected and validated for > 80% expression of CD43 by flow cytometry on two harvest days – Day 10 and Day 12. HPCs were then differentiated into iMGs over 24 days in 3F media (DMEM/F12, 2X insulin-transferrin-selenite, 2X B27, 0.5X N2, 1X Glutamax, 1X non-essential amino acids, 400 μM monothioglycerol, 5 μg/mL Insulin, supplemented with 100 ng/mL IL-34, 50 ng/mL TGFβ1, and 25 ng/mL M-CSF). Differentiated iMGs were then processed for membrane extraction or lysed and frozen at -80°C for RNA extraction. For the fourth donor, the iMG sample was derived from widely used iPSC line WTC-11 (Kreitzer et al., 2013). Differentiation was carried out by the Stem Cell Engineering Core (RRID:SCR_027503) at the Icahn School of Medicine at Mount Sinai following as described previously (McQuade et al., 2018, Matera et al., 2025).

### Immunofluorescence

Glass bottom 24-well plates (MatTek, P24G-1.5-10-F) were pre-coated with 0.01% poly-L-Lysine solution (EMD Millipore, A-005-C) for 1h at room temperature before removing and washing wells twice with cell culture grade water. Plate left uncovered and air dried in tissue culture hood for 1h at room temperature. Monocyte derived microglia (MDMi) were seeded at a density of 250,000 cells per well and cultured in RPMI media supplemented with cytokines described above. Cytokines were spiked in every other day for a period of 7 days. MDMi were fixed with 4% paraformaldehyde for 1h at 4°C. Fixed cells were washed three times with PBS. All cells were then blocked in 5% BSA and 0.1% triton X-100 in PBS for 1h at room temperature. MDMi were stained with α-IBA1 (1:500, FUJIFILM Wako Chemicals, 019-19741), α-CX3CR1 (1:500, Abcam, ab313471), and α-TMEM119 (1:200, Abcam, ab185337) in 2% BSA with 0.3% Triton X-100 overnight at 4°C. Cells were then washed 3 times for 5 mins each with 0.1% Tween-20 in PBS and α-IBA1 and α-CX3CR1 were stained with secondary Alexa Fluor-488 or FITC. α-TMEM119 staining required cells to be heated to 100°C for 4 mins in 10mM sodium citrate followed by 15 min rest at room temperature prior to blocking. Nuclear staining was done with Hoechst (1:5,000, ThermoFisher Scientific, H3570) by incubating fixed cells for 1h at room temperature prior to imaging. Stained cells were imaged 40X on Zeiss LSM980 Airyscan 2.

### RNA isolation and library preparation

Cells were collected and lysed in 100-150 µl of extraction buffer provided in PicoPure RNA isolation kit (ThermoFisher Scientific, 0204) and incubated at 42°C for 30 min shaking at 850 rpm. Lysates were stored at -80°C and later extracted according to manufacturer’s instructions. Extracted RNA was treated with a DNase-free kit (Invitrogen, 2224G) following manufacturer’s instructions. RNA was quantified using Qubit RNA High Sensitivity Assay (ThermoFisher Scientific, Q32855) and quality was assessed using Agilent RNA 6000 Nano Kit (Agilent, 5067-1511) on the Agilent Bioanalyzer 2100. Libraries were prepared using SMART-Seq Total RNA Pico Input ZapR Mammalian kit (Takara Bio, 634357) and SMART-Seq mRNA LP with UMIs (Takara Bio, 634765) according to manufacturer’s instructions. A total of 10ng of RNA was used as the starting input. For samples with low RNA concentration, input volume was maximized. Reverse transcribed cDNA was amplified with 10 PCR cycles and amplified cDNA was purified with SPRIselect beads (Beckman Coulter, B23317) following manufacturer’s size selection protocol to select for 200-700bp DNA fragments. After depletion of ribosomal cDNA, a second library amplification was performed with 18 PCR cycles and individually purified following SPRIselect beads double size selection protocol. Purified libraries were quantified with Qubit dsDNA High Sensitivity Assay (ThermoFisher Scientific, Q32854) and size were determined with Agilent High Sensitivity DNA kit (Agilent, 5067-4626). Individual libraries were normalized to 10nM and pooled together. The molarity of pooled library was determined using KAPA Library Quantification kit (KAPA Biosystems, KK4824) and measured on Azure biosystems. Pooled libraries were sequenced on the NovaSeq X Plus (Illumina) for 150bp paired-end reads.

### Membrane extraction

Cells were resuspended in ice cold homogenization buffer (250mM sucrose, 1mM EDTA, 10mM Tris HCl buffer, pH 7.5 and 2X complete protease inhibitor (Roche, 11873580001) in PBS). Cells were then homogenized on ice in a 2mL Dounce tissue grinder (KIMBLE) by 2 x 20 strokes beginning with pestle A and followed by pestle B. Samples were then sonicated with 5 x 10s pulses with 30s intervals between each pulse using the Biorupter Plus (Diagenode) cooled to 4°C with the minichiller 300 (Diagenode). Sonicated samples were centrifuged at 500xg for 10 min at 4°C to remove intact cells, nuclei and debris. Supernatants were transferred to a polycarbonate centrifuge tube (Beckman Coulter, 349622), and membrane fractions were pelleted by ultracentrifugation at 100,000xg for 1h at 4°C (Beckman Ultracentrifuge L7-65, rotor SW50.1). Pellets were washed in homogenization buffer and pelleted a second time at 100,000xg for 1h at 4°C. Pellets were then frozen at -80°C or resuspended in 150 µl of buffer (0.5% NP40, 50mM TEAB, 10mM DTT and 1x complete protease inhibitor) and frozen at -80°C until further processing.

### Sample processing for mass spectrometry

Samples for mass spectrometry were processed following a previously described protocol (Gleason et al., 2025). Briefly, precipitated samples were dissolved in 0.5% NP40, 50 mM TEAB, 10 mM DTT and sonicated. The solution was then transferred to low binding 1.5-ml vials for incubation at 70°C for 10 min with shaking. Dissolved proteins were twice precipitated using in ice-cold acetone. After acetone removal precipitates were dissolved in 100 μl of 8M urea/60 mM ABC/20 mM DTT. After 1 h of ultrasound treatment for resuspension of the pellet, samples are alkylated in 200 mM iodoacetamide. Based on the size of the pellet, the dissolved samples were split into two equal portions. This was followed by proteolytic digestion with endoproteinase LysC (Wako Chemicals) overnight and with trypsin (Promega) for 6 h at room temperature. The digestion was quenched with neat trifluoroacetic acid (Bellenguez et al.) and resulting peptide mixtures were desalted using in-house made C18 Empore (3M) StAGE tips. Approximately 1 μg of each sample was injected for analysis by reversed phase nano-LC-MS/MS (Orbitrap Ascend, Lumos, or Q Exactive separated on a 25 cm EasySprayer ThermoScientific) using 2h gradient operated in high/high.

## Bioinformatics

### RNA-seq data processing and analysis

RNAseq data in fastq format were aligned to the human reference genome (build hg38) using HISAT2 v2.2.1 with options -k 100 for a maximum of 100 alignments reported. Output BAM files were collated with SAMtools v1.17 and input into Telescope v1.0.4.1 to resolve multiple alignments among HERV loci. Raw read count tables from Telescope were normalized to trimmed mean of M values-corrected log2 counts-per-million-reads (logCPM) using the edgeR package v4.8.2 in R v4.6.1 with prior count 2 for PCA, clustering, and visualization. For both protein-coding transcriptome and HERV-specific analyses, values were filtered to include only those transcripts with at least 1 CPM in at least three samples.

### Gene expression analysis of host and HERV genes

Principal component analysis (PCA) of the transcriptome was performed with the prcomp function from the R package stats v.4.5.1 with zero-centered logCPM values for protein-coding genes (UniProtKB/Swiss-Prot, accessed 03/04/2026). Expression of a panel of 38 microglial markers across microglia models (Supplemental Table 1) was evaluated at the RNA level. PCA was also performed summarizing HERV expression with the same settings. To characterize the HERV expression profile of different microglia sources, HERVs with a median expression greater than or equal than 1 CPM within each microglia model were identified. HERVs expressed in all three model systems were plotted in a heatmap.

### Proteomics microglia source comparisons

Mass spectrometry data were searched against an amended UniProt human protein reference database (https://github.com/crgl/microglia-core-signature) using ProteomeDiscoverer/MASCOT (v.2.2). Oxidation of methionine and N-terminal protein acetylation were allowed as variable modifications, and cysteine carbamidomethylation was set as a fixed modification. An ion mass tolerance was set at 4.5 ppm for precursor and 20 ppm for fragment ions. Two missed cleavages were allowed for specific tryptic search. The ‘match between runs’ option was enabled. False discovery rates at the protein level were set to 1%. Protein abundances were represented by label free quantitation and intensity-based absolute quantitation values. To normalize results across mass spec runs, area values for each protein (grouped, 1% FDR) from Proteome Discoverer were log2 transformed and shifted so that all samples shared the same median (referred to as protein abundance). Pre-imputation transformed abundances were used for the microglia marker heatmap. Before PCA, proteins were filtered to those that were detected in at least 1/3 of samples (n=8) and missing values were imputed assuming data were missing not at random “Perseus-style”, drawn from a normal distribution downshifted 1.8 sample standard deviations from each sample mean and with a width of 0.3 sample standard deviations. Proteins were filtered to those tagged as highly confident surface in UniProt using the website API (Ahmad et al., 2025), before principal components were calculated with zero centering of normalized abundances, as described previously.

### Statistics and plotting

RNA-seq read counts were normalized based on library size using edgeR v4.10.1 and filtered to include only transcripts expressed above 1 CPM in at least three samples. For differential expression, fragment counts were transformed with the voom function of limma v3.68.4 under a model including cell source, donor sex, sequencing strandedness, and whether RNA was polyA enriched or represented the total transcriptome. To compare between groups, the average expression of genes within each cell type was compared to the average expression between the other four groups, and log2 fold changes between groups were visualized (Supplemental FIG. 1). F statistics were calculated with the empirical Bayes method with the Benjamini-Hochberg adjustment to control for multiple comparisons. Heatmaps and dendrograms were generated using the R package pheatmap v1.0.13 using complete linkage clustering and optimal leaf ordering. PCA plots were generated with ggplot2 v4.0.2 and ggprism v1.0.7 in R v4.6.1. Imputation sampling for undetected proteins was performed using NumPy v2.5.1 in Python v3.14.6.

### Gene set enrichment analysis

Gene set enrichment analysis was performed using fgsea (v1.38.0) for both RNA and proteins. For RNA, ranks for MDMi and iMG were calculated using log2 fold-changes from the one-vs-rest differential expression analysis shown in Supplemental Figure 1. For protein data, imputed abundances for surface proteins were Z-score normalized per protein, and the median value for MDMi and iMG across donors was input as ranks. GSEA was performed including the MSigDB Hallmark collection, the Gene Ontology Biological Process collection, and fetal microglia signatures from single cell datasets included in the MSigDB C8 collection. Resulting RNA assignments were filtered for adjusted p value below 0.01, and signatures for surface protein data were filtered less stringently at adjusted p value below 0.05. Pathways were collapsed, and for RNA-seq or mass spec, all signatures reaching significance for either model were visualized.

### Data and code availability

The brain-derived microglia RNA-seq dataset is a subset of the full dataset described in Kosoy et al. (Kosoy et al., 2025) and is available via the AD Knowledge portal. Donors FM_049, FM_051, FM_058, FM_093, FM_099, FM_103 were included in this study. The remainder of the RNA-seq dataset is being submitted to dbGaP (accession pending) and will be linked to the published version of this manuscript. Analyzed and raw mass spec output files are available in the PRoteomics IDEntifications Database (Project accession: accession pending).

## Results

### Comparing different microglia models

The primary sources of human microglia include brain tissue-derived microglia isolated post-mortem (BdMG), monocyte-derived microglia-like cells (MDMi), and induced pluripotent stem cell-derived microglia (iMG). Here, we compared the transcriptomic and proteomic profiles of these three microglial models using bulk RNA sequencing (RNA-seq) and mass spectrometry (mass spec), respectively. As outgroup controls, we included: primary human CD4+ T cells, NCCIT, a germ cell tumor cell line, and HMC3, an immortalized human microglial cell line.

Microglia identity is often assessed by immunofluorescence staining for limited number of canonical markers. As an illustration of this commonly used but inherently restricted approach MDMi and HeLa cells were stained Iba1 (AIF1), TMEM119, and CX3CR1. MDMi showed positive whole-cell staining for all three markers, while HeLa cells were negative although with some background present for CX3CR1 (FIG. 2a). While these results were consistent with the expression of selected microglial markers in MDMi, the approach provides limited information about overall cellular identity, heterogeneity or functional state, and was therefore not further pursued. Instead, we used an integrated transcriptomic and proteomic profiling to compare the microglial models at a systems level.

**Figure 1.**
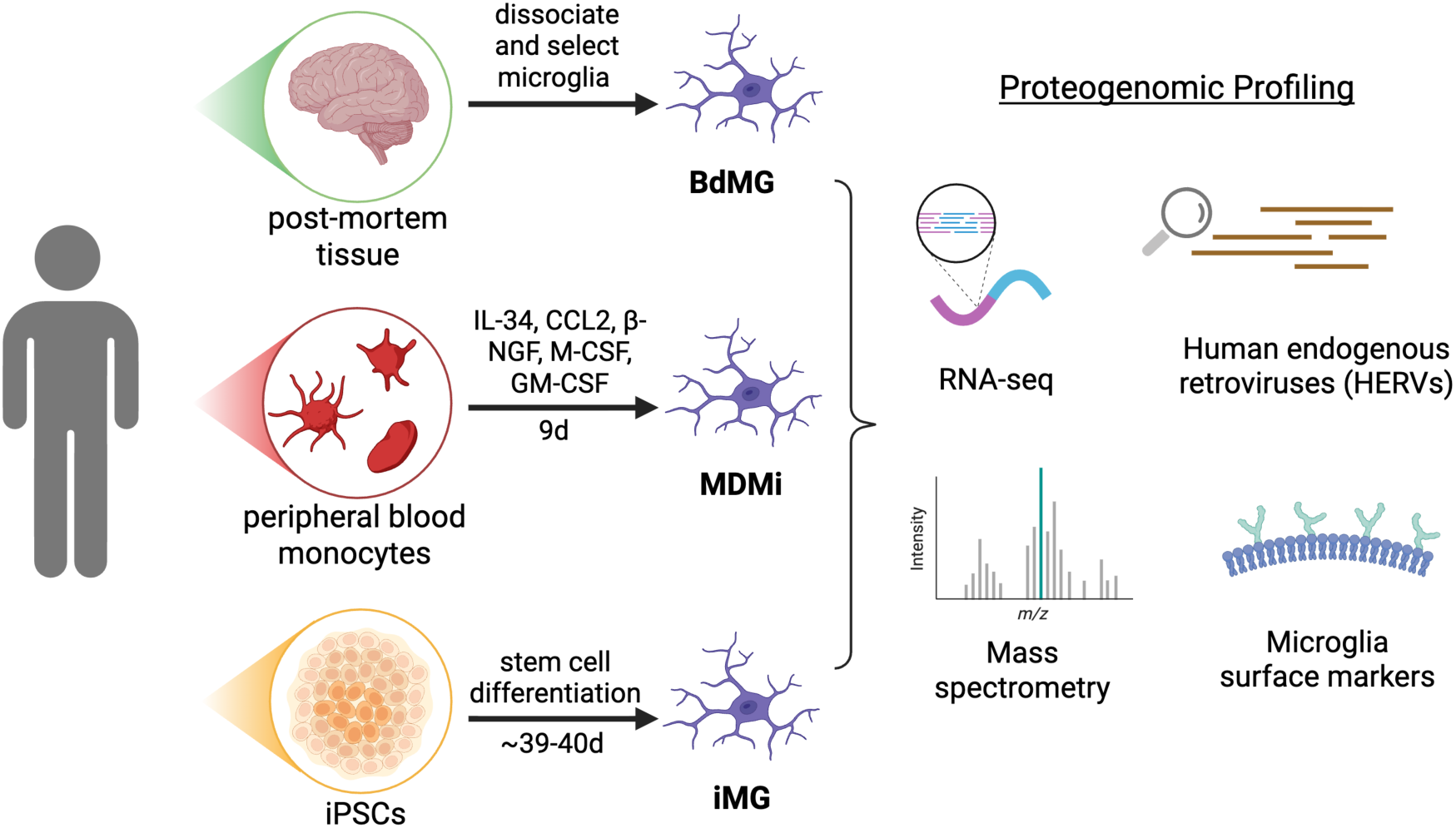
Graphical summary of the experimental design performed in this report.

**Figure 2.**
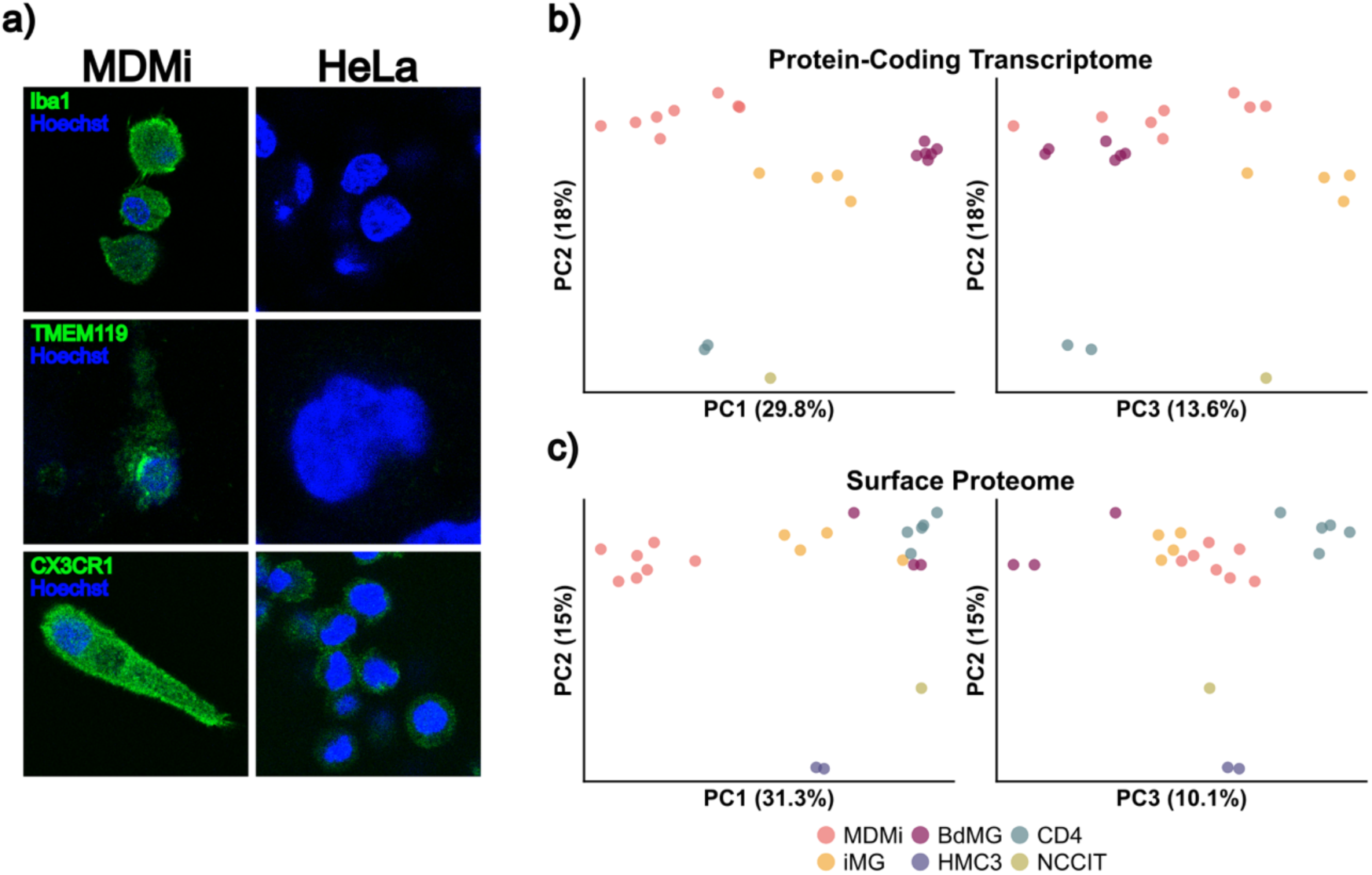
Comparison of different microglia model systems. Immunofluorescence staining of MDMi and HeLa cell control with the microglia markers, Iba1, TMEM119 and CXC3R1, and nuclear stain with Hoechst (a). Brain-derived microglia (BdMG) isolated from post-mortem tissue (n=6), monocyte-derived microglia-like cells (MDMi) (n=8), iPSC differentiated to microglia (iMG), CD4+ T-cell (n=2) and NCCIT (mean, n=3) controls were processed for RNA sequencing (RNA-seq) and cell populations visualized with principal component analysis (PCA) after filtering for protein-coding genes (b). Surface proteome of BdMG (n=3), iMG (n=4), MDMi (n=7), CD4+ T-cell (n=5), HMC3 (n=2) and NCCIT (mean, n=3) controls were identified by mass spectrometry visualized with PCA (c).

RNA-seq samples were generated from cells obtained from a total of 20 donors including 6 BdMG, 8 MDMi, 4 iMG, as well as from 3 NCCIT cell line technical replicates. Mass spectrometry was performed on 3 BdMG, 7 MDMi, 4 iMG, 5 CD4+ T cell, and 2 HMC3, and aggregated from 7 NCCIT technical replicates.

RNA-seq read counts were normalized to TMM-adjusted counts-per-million reads and log2 transformed (logCPM). Principal component analysis (PCA) of the protein-coding transcriptome revealed distinct clustering by cell type (FIG. 2b). A plot of principal component 1 (PC1) versus PC2 identified five distinct groups. PC2 separated the microglial models from the NCCIT and CD4+ T-cell outgroup controls, indicating transcriptional similarities among MDMi, iMG, and BdMG (FIG. 2a). All 3 model systems separated cleanly along PC1, while MDMi bridged BdMG and iMG along PC3, overlapping both groups. The broad dispersion of samples within the iMG and MDMi clusters likely reflects both donor-to-donor and technical variability.

PCA of surface proteins identified by mass spectrometry showed PC2 splitting the microglia model systems and CD4+ T cells from HMC3 and NCCIT cell lines (FIG. 2c). PC1 separates out MDMi from all other cell types tested with broad dispersion within and across groups, while PC3 clusters the MDMi and iMG model systems between CD4+ T cells and BdMGs. The dispersion of samples within each group suggests that donor-related and technical variability is also evident at the protein level.

Together, these initial findings indicate that MDMi share transcriptomic and proteomic properties with human microglia, although, to provide information for the identification of additional microglial markers and about the activation states represented by the different model systems, a more detailed analyses of the RNA-seq and mass spectrometry data is necessary.

### Microglia models share common surface marker RNA expression profile

A panel of 38 representative microglia surface markers identified from the literature (Table S1) (Bellenguez et al., 2022, Martins-Ferreira et al., 2025, Keren-Shaul et al., 2017, Butovsky et al., 2014, Esaulova et al., 2020, Olah et al., 2020, Masuda et al., 2019, Galatro et al., 2017, Hickman et al., 2013) was used to evaluate the microglia phenotype. A subset of 36 out of the 38 surface markers had detectable expression in our dataset and were visualized on a heatmap comparing microglia models to CD4+ T cell and NCCIT cell negative controls (FIG. 3a). Unsupervised clustering was performed, and samples clustered amongst their respective groups, indicating similarities within each sample type irrespective of donor. As expected, NCCIT and CD4+ T cell controls clustered separately from MDMi, iMG and BdMG (FIG. 3a).

**Figure 3.**
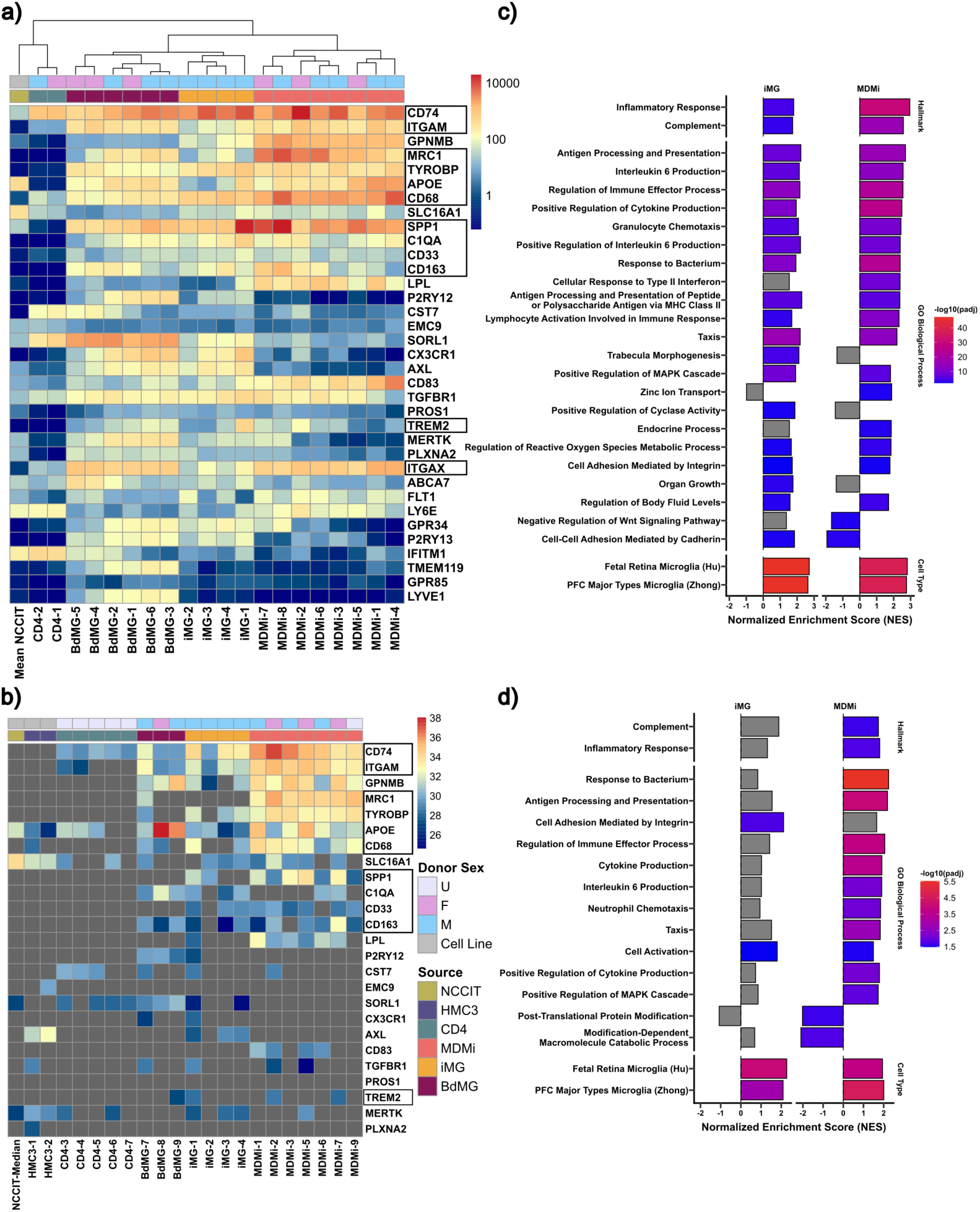
A consensus surface marker profile in microglia. RNA transcripts and proteins in microglia samples or controls matched to 36 (RNA) and 24 (protein) of 38 surface markers. Unsupervised clustered heatmap of surface marker genes in each sample and cell type: brain-derived microglia (BdMG, n=6), iPSC-derived microglia (iMG, n=4), monocyte-derived microglia (MDMi, n=8), CD4+ T-cell (n=2) and NCCIT (mean, n=3) controls (a). Mass spectrometry proteins identified in microglia, BdMG (n=3), iMG (n=4) and MDMi (n=7) and non-primary microglia controls, HMC3 (n=2), CD4+ T-cell (n=5) and NCCIT (median, n=10, biological replicates) matched to surface proteins in RNA-seq (b). Proteins not detected by mass spec in a given sample are represented with grey boxes. Samples are annotated with sex and cell type. Colors correspond to CPM (a) and abundance (b). GSEA results are plotted for RNA-seq (c) and mass spec (d). Gene sets are grouped by category and sorted within category by maximum significant normalized enrichment score across iMG and MDMi, with colors indicating adjusted p value. Color scales are independent for each panel. Gene sets that do not reach significance (p>0.01 and p>0.05 for RNA and protein data, respectively) are colored grey.

Expression of each gene was averaged across sample type, and microglia were compared to CD4+ T cell or NCCIT controls. We identified a subset of genes expressed consistently in all microglia models (geometric mean CPM of at least 10 in all models) which were less highly expressed (>10-fold lower) in at least one of NCCIT or CD4 controls (*CD74*, *ITGAM*, *GPNMB*, *MRC1*, *TYROBP*, *APOE*, *CD68*, *SPP1*, *C1QA*, *CD33*, *CD163*, *TREM2*, *ITGAX*, FIG. 3a, boxes). Of these, *CD74, TYROBP*, *C1QA*, and *CD33* were enriched to a similar degree (within 4-fold) in all microglia models (Supplemental FIG. 1). The microglia signature genes *GPNMB* and *MRC1* were highly variable across model systems and particularly enriched in MDMi (Supplemental FIG. 1). Some of these genes, including *APOE*, *C1QA*, *TYROBP*, and *TREM2*, contribute to microglia-mediated immune responses and have been implicated in Alzheimer’s disease (Fonseca et al., 2017, Haure-Mirande et al., 2017, Konishi and Kiyama, 2018). APOE, which was upregulated in BdMG and MDMi, has three main variants (ε 2, ε 3, ε 4) that significantly alter the risk for late-onset Alzheimer’s disease (Corder et al., 1993, Krasemann et al., 2017, Farrer et al., 1997, Keren-Shaul et al., 2017, Kosoy et al., 2025). Both TREM2 and TYROBP, found upregulated in all three microglia model systems, form a cell-surface signaling complex that regulates microglial survival, proliferation, and phagocytosis, as well as the transition from a homeostatic phenotype toward a disease-associated microglial (DAM) state (Audrain et al., 2021, Linnartz-Gerlach et al., 2019, Konishi and Kiyama, 2018).

We also detected transcription of genes encoding canonical microglia markers, *P2RY13*, *TMEM119, CX3CR1, ITGAM* (*CD11b*)*, MRC1* (*CD206*)*, CD68* and *ITGAX* (*CD11c*) (FIG. 3a), although their expression varied among the three models. Average expression of *P2RY13* and *CX3CR1* was more than 50-fold lower in MDMi compared to iMG and BdMG, while average expression of *TMEM119* was more than 80-fold lower in MDMi and iMG compared to BdMG. One possible explanation for this phenotype is that these microglia markers are heavily regulated by cell activation status. Previous studies have shown that DAM exhibit reduced expression of homeostatic markers, including TMEM119, CX3CR1, and P2RY13, accompanied by increased expression of activation and disease-associated genes such as *APOE*, *C1QA*, *TYROBP*, *TREM2*, *ITGAX*, *CD68*, and *SPP1* (Keren-Shaul et al., 2017, Krasemann et al., 2017, Böttcher et al., 2019, Kenkhuis et al., 2022). Increased expression of MRC1 is also observed in M2-like microglia activation (von Ehr et al., 2020). Taken together, these expression patterns suggest that BdMG retain a comparatively homeostatic profile, whereas MDMi and iMG display more activation-associated features. Nevertheless, the shared expression of multiple microglial identity and functional markers across the three groups demonstrates substantial molecular similarity among these model systems.

### Surface marker protein expression profiles identified across microglia model systems

Although surface marker transcripts were detected by RNA-seq, the correlation between steady state mRNA and protein abundance is weak (Koussounadis et al., 2015) as translation is regulated by additional cellular mechanisms (Payne, 2015, Schwanhäusser et al., 2011, Liu et al., 2016, Vogel and Marcotte, 2012). Therefore, we sought to characterize the surface protein profile of our microglia models. Membrane-enriched fractions of microglia model systems and outgroup control cell types were isolated and analyzed by mass spec. The membrane-enriched mass spec data was filtered against the same subset of 38 microglial marker proteins, with 22 of a possible 38 proteins appearing in at least one microglia sample (FIG. 3b). All 24 markers (22 in primary microglia and 2 only in HMC3) detected were also among the 36 detected at an RNA level. Comparison of RNA-seq and mass spec data identified 9 overlapping markers appearing in at least one donor within all 3 microglia model systems (CD74, ITGAM, GPNMB, MRC1, TYROBP, APOE, CD68, SPP1, C1QA, CD33, CD163, and TREM2 FIG. 3b, boxes). Membrane-associated proteins ITGAM (CD11b), TYROBP, and CD68 were either low or undetectable in NCCIT, CD4+ T cell and HMC3 controls compared to BdMG, iMG and MDMi (FIG. 3b). Notably, CD33, a protein typically found in myeloid and monocytic lineage immune cells (De Propris et al., 2011), was also detected in MDMi and iMG (FIG. 3b), revealing similarities between the two model systems. Other markers, including MRC1, SPP1, and LPL, were similarly detected in iMG and MDMi samples but not in BdMG.

Microglia markers CX3CR1 and P2RY12 proteins were detected only in a subset of BdMG and iMG donors (FIG. 3b). There were 13 surface markers detectable by RNA-seq but not mass spec (FIG. 3a-b), likely due to a combination of the lower sensitivity of mass spec compared to RNA-seq and the highly variable relationship between steady state RNA and protein levels across genes. Indeed, we observed *TMEM119* expression at the transcriptome level (FIG. 3a), as well as at the protein level using immunofluorescence staining of MDMi (FIG. 2c), but mass spec failed to detect TMEM119, confirming limitations of mass spec-mediated detection used in isolation.

### Increased immune signaling and microglia activation in MDMi and iMG

We performed Gene Set Enrichment Analysis (GSEA) to match RNA and proteins identified in iMG and MDMi to hallmark genes, biological processes and cell types. The Molecular Signatures Database (MSigDB) hallmark gene sets and Gene Ontology (GO) Biological Process and fetal microglia signatures from the MSigDB cell type gene sets were analyzed (Subramanian et al., 2005, Liberzon et al., 2011, Liberzon et al., 2015). GSEA revealed iMG and MDMi were significantly enriched for genes belonging to inflammatory response and complement cascade hallmarks (FIG. 3c). When transcripts were matched to GO biological processes, immune response pathways were significantly enriched in both iMG and MDMi (FIG. 3c).

At the protein level, the same hallmark gene set (inflammatory response and complement cascade) was significantly enriched in MDMi samples, but not in iMG (FIG. 3d). Of the biological processes identified in mass spec, more pathways were significantly enriched in MDMi compared to iMG (10 and 2 respectively) (FIG. 3d). From a functional standpoint, these findings suggest that MDMi retain more immune functions compared to iMG and behave more like activated microglia. This difference was most striking at a protein level, where iMGs show a significant enrichment of the fetal microglia signature but enrichment of immune signatures does not reach the level of significance. This further supports MDMi as a practical and effective alternative *in vitro* model for studying microglia.

### Defining a HERV transcriptome signature in microglia

Having established that these models share a core surface marker signature and recapitulate an immune-activation phenotype, we next asked whether their similarity extends to a resident retroviral axis directly relevant to neuroinflammation. HERVs are challenging to study due to repetitive nature of their sequences and low mRNA and protein expression (Bowles et al., 2022). Previous reports have shown that HERV expression is likely tissue and cell-type specific (She et al., 2022, Zhu et al., 2025). Some HERV families have been shown to be upregulated and associated with microglia-mediated proinflammatory signaling in AD and MS (Dembny et al., 2020, Adler et al., 2024, Duarte et al., 2025). This underscores the necessity to characterize HERVs in different cell types to better understand their role in pathogenesis. Although transcriptomics studies have been performed investigating HERVs in microglia, HERVs in MDMi and iMG remain understudied. To identify the HERV transcriptome with locus-level resolution in our microglia model systems, we employed Telescope, a specialized bioinformatics tool used to accurately estimate transposable element expression from RNA-seq data (Bendall et al., 2019). PCA on the HERV transcriptome revealed clustering within cell types (FIG. 4a). Analyzing only HERV expression, PC1 separates BdMG from other microglia models and controls, while PC2 pulls the NCCIT cell line away from primary cell samples.

**Figure 4.**
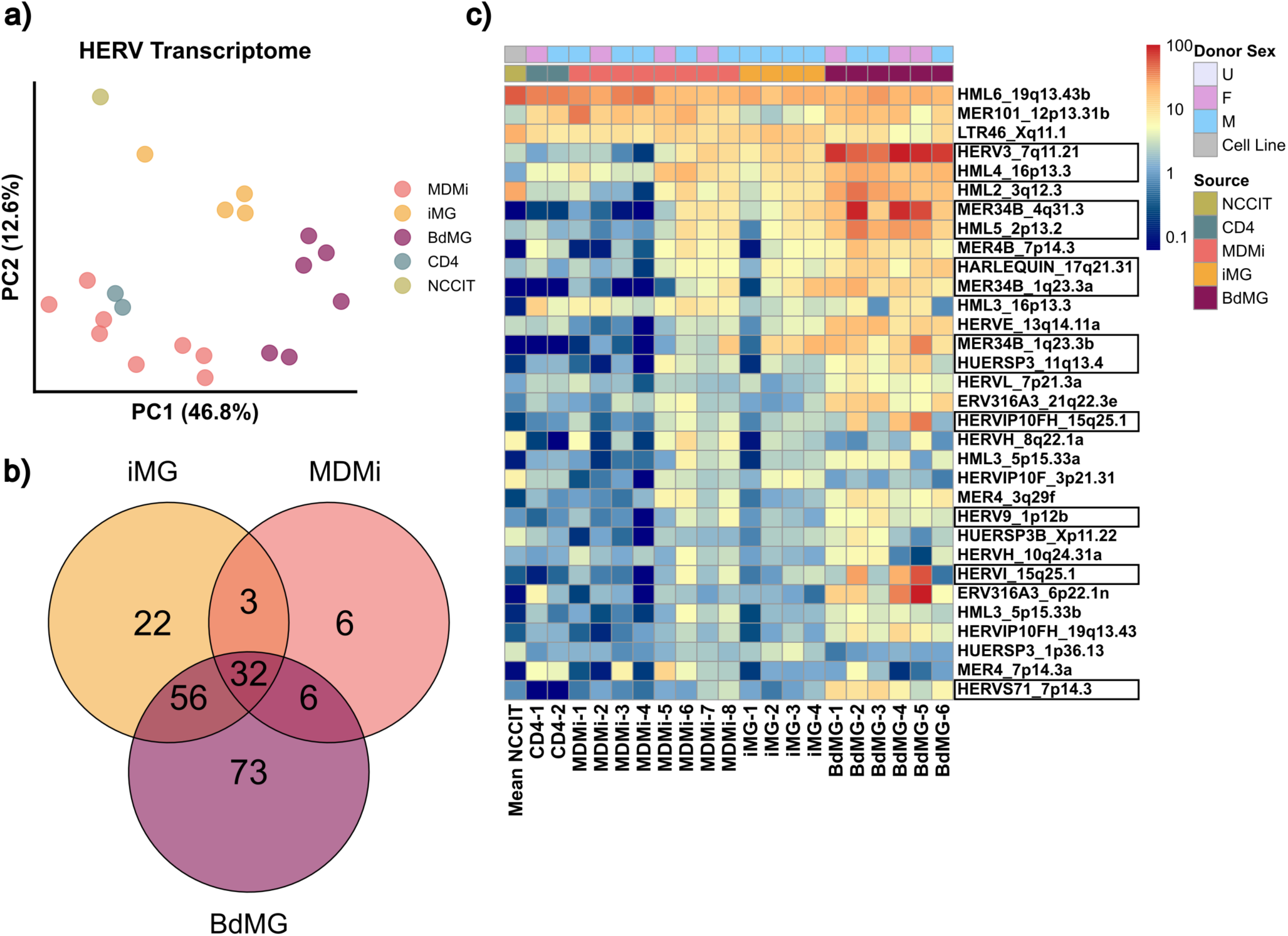
A consensus HERV profile in microglia. Principal component analysis (PCA) of brain-derived microglia (BdMG, n=6), iPSC-derived microglia (iMG, n=4), monocyte-derived microglia (MDMi, n=8), CD4+ T cell (n=2) and NCCIT (geometric mean of 3 technical replicates) (a) and comparison of number of unique or overlapping HERV transcripts with at least one CPM identified across microglia models (b). Heatmap representing the 32 overlapping HERV transcripts identified with color keys represent either donor sex or cell source for each gene across samples ranked by descending median expression across all samples (c). Boxes represent HERVs with maximum expression in each microglia model system at least 2-fold higher than maximum expression in controls.

To identify commonly expressed HERV transcripts within each microglia model system, retroviral elements were filtered to include those with a median expression of at least 1 CPM. We identified 47 HERVs in MDMi, 113 HERVs in iMG and 167 in BdMi (FIG. 4b). Of those, 32 HERVs were present in all three microglia models (FIG. 4b). Unclustered heatmap analysis of the 32 overlapping HERVs showed several HERVs enriched in microglia models (HERV3_7q11.21, HML4_16p13.3, MER34B_4q31.3, HML5_2p13.2, HARLEQUIN_17q21.31, MER34B_1q23.3a, MER34B_1q23.3b, HUERSP3_11q13.4, HERVIP10FH_15q25.1, HERV9_1p12b, HERVI_15q25.1, HERVS71_7p14.3, FIG. 4c, boxes) compared to negative controls. Specifically, HERVs were considered enriched if the maximum expression across donors for all model systems was at least 2-fold greater than the maximum expression across negative controls. Donors MDMi 1-4 and iMG-1 displayed lower HERV expression compared to other MDMi and iMG samples, which can also be observed in the PCA plot (FIG. 4a), highlighting real life donor differences as opposed to the homogeneity found when assessing transformed cell lines.

The HERV transcriptome profile is also composed of different clade members, implying that no singular clade is dominant in microglia. Of the HERVs identified, some have been previously implicated in different diseases. Transcripts of HERV-K HML4 have been detected in post-mortem tissues of AD patients (Dawson et al., 2023). Here we see a common HERV transcriptome profile across all microglia models, suggesting that HERV expression in microglia is conserved at steady state.

## Discussion

Microglia play essential roles in brain immunosurveillance and maintenance of central nervous system (CNS) homeostasis (McKinney et al., 2026, Colonna and Butovsky, 2017). As the primary responders to CNS injury, they eliminate pathogens, dead cells, and other potentially harmful antigens that threaten CNS integrity (Colonna and Butovsky, 2017, Sargeant and Fourrier, 2023, Cuadros et al., 2022). Microglia also regulate neuroinflammatory responses and can adopt either pro-inflammatory or anti-inflammatory phenotypes depending on the nature of the stimulus (Colonna and Butovsky, 2017, Smith et al., 2012). Reports have shown that microglia may protect against neurodegenerative disease by phagocytosing misfolded proteins; however, they can also contribute to disease progression by releasing inflammatory cytokines that enhance protein aggregation and neuronal injury (Gao et al., 2023, Baaklini et al., 2023, van Wageningen et al., 2019, Noh et al., 2025, Bond et al., 2025). Common *in vitro* approaches for studying microglia include immortalized cell lines, induced pluripotent stem cell-derived microglia (iMG), monocyte-derived microglia-like cells (MDMi), and microglia isolated from post-mortem brain tissue (Gumbs et al., 2022, Woolf et al., 2025, Ryan et al., 2017). The HMC3 immortalized cell line is a convenient model but, based on our findings, their profile differs widely from those of BdMG, iMG, and MDMi, making their physiologic relevance uncertain (FIG. 3b). Conversely, BdMG are the most physiologically relevant, but access to post-mortem brain tissue is limited and yields are often low with poor viability. iMG are more accessible but their differentiation requires a lengthy protocol of approximately 40 days. Finally, MDMi offer a practical model that overcomes several of these limitations while preserving physiological relevance. In this study, we integrate transcriptomics and proteomics to identify a common profile across microglia model systems that establishes their similarities both at the RNA and protein level.

In our transcriptome analysis, we identified overlapping microglia surface markers across most of our models. We also detected expression of microglia activation markers, APOE, C1QA, TYROBP, TREM2, ITGAX, MRC1, CD68 and SPP1 (FIG. 3a). Many of these include immune-response markers and have been associated with Alzheimer’s disease (AD). Apolipoprotein E (APOE) is known to colocalize and influence metabolism of amyloid-β (Aβ) plaques, a hallmark of AD (Namba et al., 1991, Wisniewski and Frangione, 1992, Verghese et al., 2013). APOE also influences microglia activation, potentially regulating the innate immune response to Aβ-plaques (Corder et al., 1993, Krasemann et al., 2017, Farrer et al., 1997, Keren-Shaul et al., 2017, Ulrich et al., 2018, Kosoy et al., 2025). TREM2 is primarily expressed by microglia in the CNS and associates with the adaptor protein, TYROBP (DAP12), during microglia activation (Hickman et al., 2013, Atagi et al., 2015). Interestingly, APOE also binds TREM2 and may be involved with phagocytosis (Atagi et al., 2015). Other surface proteins, CX3CR1, ITGAM (CD11b) and ITGAX (CD11c) help control migration and positioning of microglia in the CNS, with ITGAX also known to be upregulated in activated microglia (Colonna and Butovsky, 2017). Complement component 1q (C1q) is produced and secreted by microglia to mediate synaptic pruning and stripping (Scott-Hewitt et al., 2024, Colonna and Butovsky, 2017). C1qA belongs to one of six heterotrimers in the C1q complex (Mortensen et al., 2017) and is observed across all microglia models by RNA-seq (FIG. 3a). Secreted phosphoprotein 1 (SPP1) is required for activation of C1q and both together shift microglia to a phagocytic state (De Schepper et al., 2023). SPP1 has also been found in plaque-associated microglia in AD patients (De Schepper et al., 2023).

When assessing protein-level expression of microglia markers, we detected 22 surface markers, with each appearing in at least one donor for at least one microglia model (excluding the HMC3 cell line) (FIG. 3b). The failure to detect some proteins detected at the RNA level may reflect limited sample input and/or insufficient mass spec sensitivity. Mass spec has known limitations, including reduced sensitivity for low-abundance proteins and susceptibility to sample complexity (Fonslow et al., 2011, Wacholder and Carvunis, 2023). Nonetheless, even though proteins may go undetected, a positive mass spec signal provides strong evidence that the protein is expressed in the sample. These findings underscore the limited relationship between RNA and protein expression, supporting the use of complementary approaches when identifying proteins of interest. Indeed proteins, rather than RNA transcripts, are typically the functional effector molecules, therefore integrating transcriptomic and proteomic data provides a more informative understanding of microglia identity. By taking a proteogenomic approach, we identify a shared microglia marker profile at both the transcript and protein-level for the different microglia models (FIG. 3a-b) and reinforce the importance of using a multi-marker panel for identifying microglia populations instead of relying on individual markers.

Performing GSEA on both the MDMi and iMG transcriptomes and proteomes allows for a high-level comparison of the RNA-seq and the protein expression profiles. We observed significant enrichment of immune pathways in both MDMi and iMG by RNA-seq, but this was not the case for mass spec (FIG. 3c-d). While MDMi retained significant enrichment of immune pathways, iMG hallmark and biological processes that were significantly enriched in RNA-seq were not significant at the protein level (FIG. 3d). Only cell activation was significantly enriched in both iMG and MDMi (FIG. 3d), corroborating our observation that MDMi and iMG are in an activated state.

This is likely due to the different lineages of the two models. MDMi are derived from peripheral blood monocytes, which are myeloid cells that originate from hematopoietic stem cells in the bone marrow (Kawamoto and Minato, 2004, Zhao et al., 2012, Ziegler-Heitbrock, 2015), while iMG are differentiated from induced pluripotent stem cells (iPSCs) that are reprogrammed adult somatic cells isolated from skin or blood (Yu et al., 2007, Park et al., 2008, Takahashi and Yamanaka, 2006). Nonetheless, many surface markers are consistently identified across all microglia models, making MDMi and iMG comparable models to BdMG.

HERVs are typically dormant under homeostatic conditions but have been increasingly linked to the pathogenesis of neurodegenerative diseases. In MS, HERV-W RNA has been detected in retrovirus-like particles, as well as in patients’ brain lesions, cerebrospinal fluid, lesions, and peripheral blood mononuclear (Perron et al., 1989, Komurian-Pradel et al., 1999, Dolei et al., 2002, Perron et al., 2012, van Horssen et al., 2016, Perron et al., 1997, Mameli et al., 2007, Rasmussen et al., 1995). HERV-K include 11 distinct clades (HML1-11) (Subramanian et al., 2011). HERV-K transcripts have been identified in serum from patients with ALS (Phan et al., 2021). In AD, transcriptomic studies have revealed elevated HERV-W, HERV-K HML2, HML3, HML4, and HML6 expression compared to health controls in tissue derived from postmortem brains (Dawson et al., 2023).

Using Telescope to quantify transcripts aligning to retroviral loci, we identify a HERV transcript profile across different microglia models (FIG. 4c), including HERV-K members that are proximal to host genes such as oncogenic marker KCTD5 (HML4_16p13.3) (Shi et al., 2023) and transcription regulator zinc finger protein ZNF638 (HML5_2p13.2) which represses the transcription of HBV closed circular DNA. Furthermore, HML2_3q12.3 is one of the most broadly expressed HML-2 proviruses in healthy human tissues (Burn et al., 2022) and is reported to encode an intact gag ORF with protein production potential (Srinivasachar Badarinarayan and Sauter, 2022). Beyond the HERV-K clade, HERV3_7q11.21 appeared to be highly expressed in brain-derived microglia (FIG. 4c). It encodes the intact surface domain (SU) of the envelope protein, and is reported to be expressed in the placenta (Andersson et al., 2005), and other tissues undergoing fusion such a skeletal muscle and tumor associated macrophages (Grandi and Tramontano, 2018).

These findings suggest that specific HERV loci may be transcriptionally active in microglial cells and could contribute to aspects of microglial identity, maintenance, or homeostatic function. Although the precise biological roles of these elements remain undefined, their consistent detection raises the possibility that HERV expression is not merely transcriptional noise but may reflect regulated activity linked to microglial state or function. These elements may indeed be components of regulatory networks that modulate innate immune signaling, inflammatory responsiveness, cellular maturation or adaptation to environmental cues. Future studies examining locus-specific regulation, chromatin accessibility, response to inflammatory stimulation, and effects of HERV perturbation will be important to determine whether these elements actively shape microglial phenotypes.

Some limitations of this study include donor differences and batch effects from different RNA-seq and tissue processing. BdMG RNA-seq was performed separately, and RNA-seq batch effects could not be corrected. In addition, the sensitivity of mass spectrometry remained a limiting factor for comprehensive detection of low-abundance proteins indicating that an integrated proteogenomic approach (transcriptomic plus Mass Spec) is more effective for a accurate microglia cell type identification. Finally, it is evident that microglia marker expression in MDMi and iMG can be influenced by differentiation protocols and culture conditions as opposed to *in vivo* microglia (Melief et al., 2016, Jantti et al., 2024). However, few studies have directly compared all three model systems at both the transcript and protein-level. Here we show that MDMi and iMG exhibit similar phenotypes relative to BdMG, with MDMi displaying greater enrichment in immune functions compared to iMG. Based on these findings, we conclude that MDMi are the most efficient, accessible and practical model for microglia studies.

Additionally, the use of MDMi as a model for microglia physiology underscores the practical and translational value of collecting readily accessible patient material. Peripheral blood monocytes, obtained through minimally invasive sampling, can be differentiated to MDMi, permitting researchers to longitudinally assess changes in microglial-like function such as inflammatory responses, phagocytic capacity, as disease progresses or in response to therapeutic intervention. This has recently acquired even more relevance since the publication of a report that indicates that, through human lifespan, brain microglia are replaced with microglia originating from the same myeloid precursors, monocytes derive from (Belk et al., 2026).

## Acknowledgements

Microscopy and image analysis was performed at the Microscopy and Advanced Bioimaging CoRE at the Icahn School of Medicine at Mount Sinai (RRID: SCR_027237). We would like to thank the Mount Sinai Neuropathology Brain Bank & Research CoRE (RRID: SCR_027565) for the collection of autopsy brain specimens used in this study.

We would like to thank the staff of the Stem Cell Core facility, notably Hemanta Sarmah, PhD, under the leadership of Barbara Corneo, PhD, at Columbia University Irving Medical Center for their contributions to the work presented in this manuscript. We would also like to thank the Stem Cell Engineering Core (RRID:SCR_027503) at the Icahn School of Medicine at Mount Sinai for providing a portion of the iMGs used in this study.

## Contributions: CREDIT

**H.T.W.** Conceptualization, Mass spectrometry, data analysis, Methodology, Investigation, Writing, Review and editing.

**C.G.** Conceptualization, Pipeline development, Sequencing analysis, Mass spectrometry data analysis, Visualization, Review and editing.

**A.B.** Pipeline development, Sequencing analysis, Mass spectrometry data analysis, Visualization, Review and editing

**M.A.** Investigation, Review and editing.

**R.K.** Sequencing analysis.

**J.F.F.** Investigation, Review and editing.

**E.L.** Methodology, Investigation.

**G.E.H.** Sequencing analysis.

**H.M.** Mass spectrometry, Mass spectrometry data analysis.

**P.R.** Conceptualization, Funding Acquisition

**D.B.** Conceptualization, Funding Acquisition

**V.S.** Conceptualization, Funding Acquisition, Review and editing

**L.C.F.M.** Conceptualization, Methodology, Visualization, Funding Acquisition, Writing, Review and editing.

---

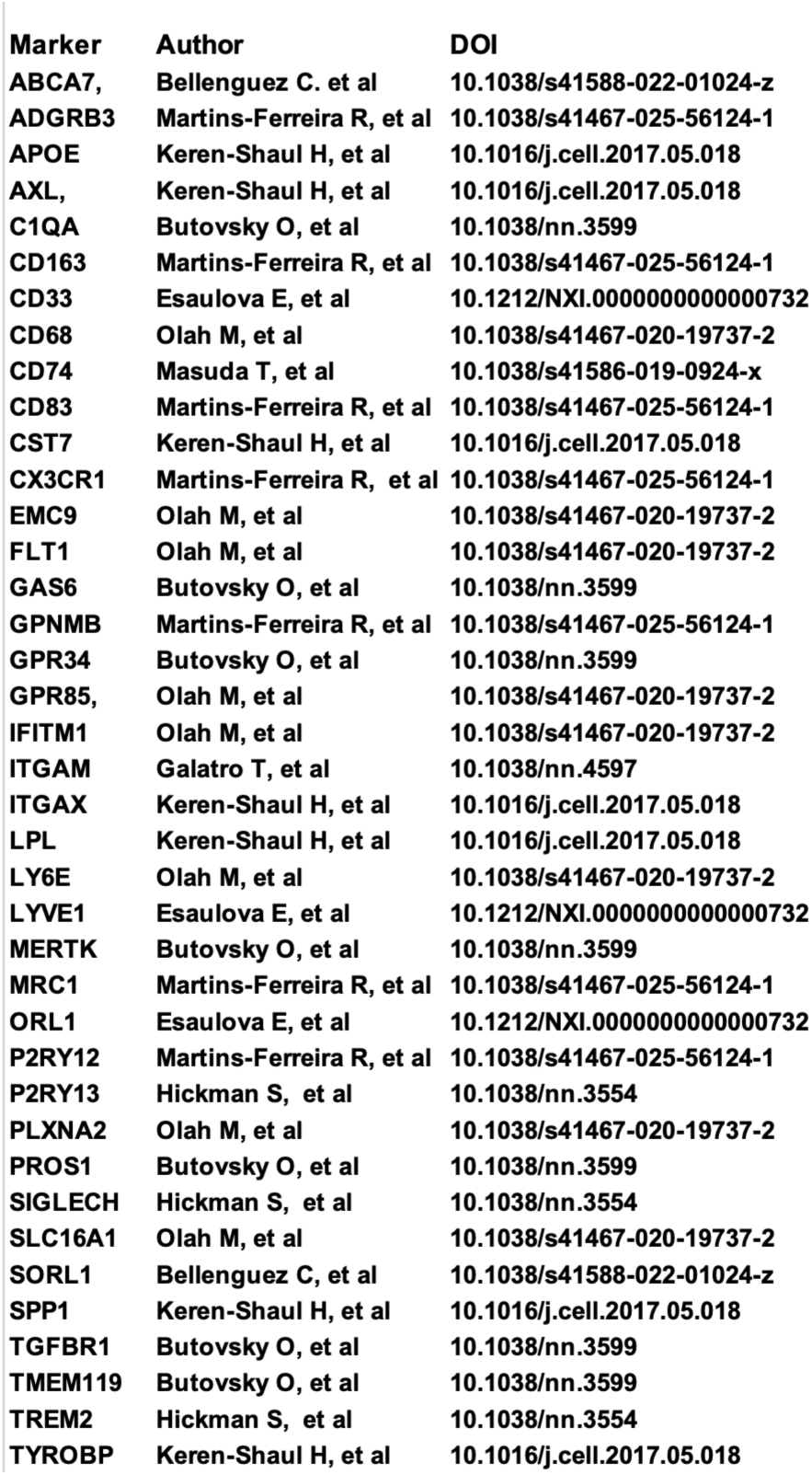
Supplemental Table 1.

**Supplemental Figure 1.**
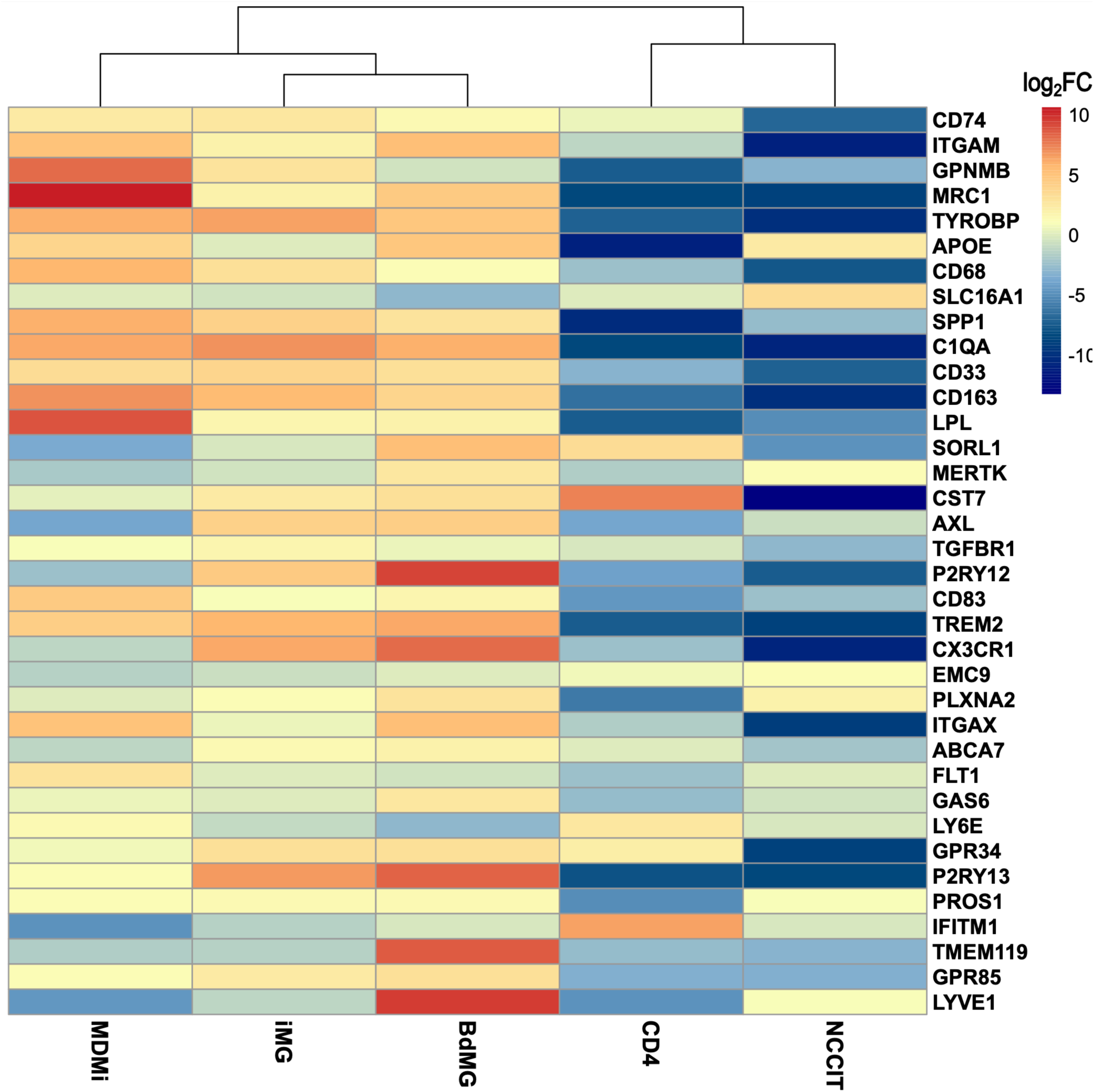
Log2 fold change of expression of microglia signature genes compared to outgroup means between model systems MDMi (n=8), iMG (n=4), and BdMG (n=6), and CD4+ t-cell (n=2), and NCCIT (n=3, technical replicates) negative controls.

